# Long-term breast cancer response to CDK4/6 inhibition defined by TP53-mediated geroconversion

**DOI:** 10.1101/2023.08.25.554716

**Authors:** Rei Kudo, Anton Safonov, Edaise da Silva, Qing Li, Hong Shao, Marie Will, Atsushi Fushimi, Harikrishna Nakshatri, Jorge S. Reis-Filho, Shom Goel, Andrew Koff, Britta Weigelt, Qamar J. Khan, Pedram Razavi, Sarat Chandarlapaty

## Abstract

Inhibition of CDK4/6 kinases has led to improved outcomes in breast cancer. Nevertheless, only a minority of patients experience long-term disease control. Using a clinically-annotated cohort of patients with metastatic HR+ breast cancer, we identified *TP53* loss (28.8%) and *MDM2* amplification (6.7%) to be associated with lack of long-term disease control. Human breast cancer models revealed that p53 loss did not affect CDK4/6 activity or G1-blockade, but instead promoted drug-insensitive p130 phosphorylation by CDK2. Persistence of phospho-p130 prevented DREAM complex assembly, enabling cell cycle reentry and tumor progression. Inhibitors of CDK2 could overcome p53 loss, leading to geroconversion and manifestation of senescence phenotypes. Complete inhibition of both CDK4/6 and CDK2 kinases appears to be necessary to facilitate long-term response across genomically-diverse HR+ breast cancers.

## Introduction

Inhibition of the G1 checkpoint is an integral requirement for numerous oncologic therapies and has been most effectively realized in hormone receptor positive (HR+) breast cancer through the use of selective CDK4/6 kinase inhibitors (CDK4/6i) (*1, 2*). Cells held in G1 by CDK4/6 inhibition can undergo a variety of fates, including apoptosis, senescence, or cell cycle reentry (*3, 4*). Of those, cell cycle re-entry is a major limitation to the efficacy of these drugs as it can restore proliferative capacity and allow mutagenesis that can further enforce resistance. By contrast, eliciting a cell state that prevents cell cycle re-entry can promote tumor clearance and long-term disease control (*5*).

In HR+ breast cancer, CDK4/6i are standard of care promoting relapse free and overall survival (*6–11*). However, there is high variance in the degree and duration of benefit from these therapies ranging from a few months to over 5 years (*12–14*). The initial effect of the CDK4/6i is to cause a marked decline in cell cycle genes in the vast majority of tumors, closely corresponding to their induction of G1 arrest in laboratory models (*15*) (*16*). However, a rebound in Ki67 levels is observed during prolonged therapy, suggesting that some tumors have the ability to escape drug-mediated G1 arrest (*17*) (*18*). We hypothesized that this capacity for cell cycle re-entry might cause the lack of long-term disease control.

In this study, we sought to determine the genomic configurations and underlying mechanisms associated with long-term disease control among patients on CDK4/6i and thereby identify pharmacologic approaches to elicit this effect in the majority of patients.

## Results

### Somatic variants enriched in cancers failing to have long−term response to CDK4/6i

To identify genomic patterns associated with clinical outcomes, we analyzed a cohort of 447 patients with metastatic hormone receptor-positive and HER2-negative (HR+/HER2–) breast cancer treated at MSK with first-line CDK4/6i (abemaciclib, palbociclib, ribociclib), tumor-normal sequencing of up to 468 cancer-associated genes was performed via the FDA-authorized MSK-IMPACT assay (*20*). For each gene, we defined somatic mutations, structural variants, copy-number alterations and select germline pathogenic variants as identified by OncoKB (*21*). First, we sought to identify alterations in individual genes associated with longer, intermediate and short response. This was accomplished by dividing patients by time on first-line CDK4/6i and ET into tertiles (consisting of 149 patients each). Patients who experienced intermediate or short response and discontinued therapy for reasons other than disease progression were excluded from the analysis. We iteratively assessed enrichment of oncogenic variants in each gene with a Firth-penalized logistic regression, comparing cases with longer response to i) intermediate response and ii) short response. The model was adjusted for de novo metastatic status at diagnosis and endocrine therapy partner, and corrected for multiple hypothesis testing. Oncogenic alterations in *PPM1D* (odds ratio [OR]: 3.64, 95% confidence interval [CI]: 1.45-9.15, p = 0.006, q = 0.09) and *TP53* (OR: 2.16, 95% CI: 1.23-3.78, p = 0.007, q = 0.09) were significantly enriched in the intermediate, compared to long-term response group after adjustment for multiple-hypothesis testing (Fig.1A). In patients who experienced a short response, *MDM2* (OR: 3.26, 95%CI: 1.10-9.67, p = 0.032, q = 0.26)*, MYC* (OR: 2.75, 95%CI: 1.22-6.16, p = 0.015, q = 0.17) and *TP53* (OR: 4.43, 95%CI: 2.45-8.02, p < 0.001, q < 0.001) mutations were putatively enriched in the short response group. *TP53* alterations remained significant after adjustment for multiple-hypothesis testing (Fig 1B).

**Figure 1.**
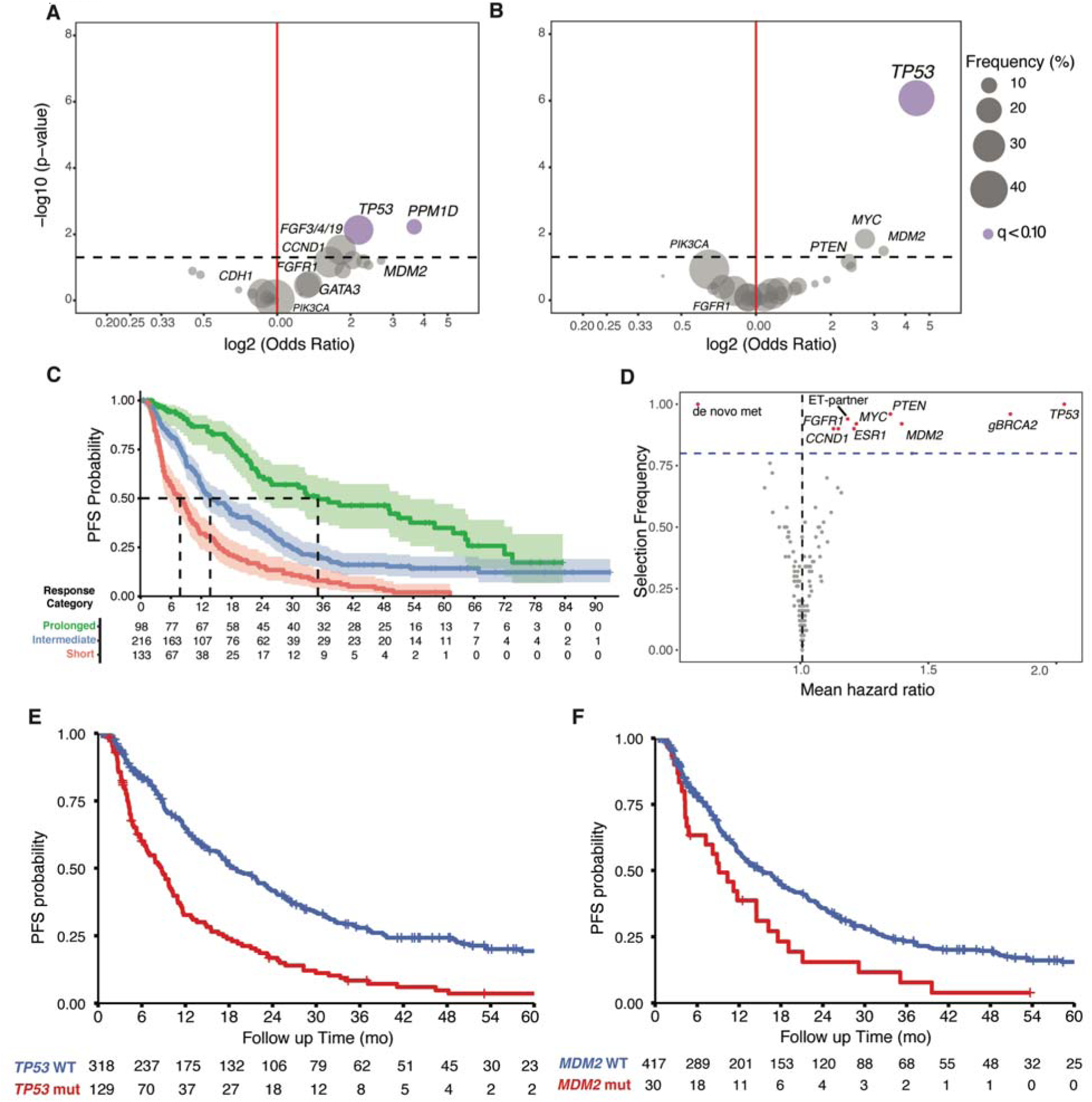
Somatic variants enriched in patients failing to achieve long−term response to CDK4/6i. The association of individual genes with long term disease control (PFS) as compared to **A) inter-mediate and B) short-term disease control**, CDK4/6i and ET, based on even tertiles of time on treatment is depicted. Color indicates statistical significance by (q < 0.10) and size of the circle reflects the frequency of alteration in the cohort. All q values are calculated based on Benjamini and Hochberg method correction of log-rank p values. **C,** depicts the results of an elastic net Cox proportional hazard model, whereby risk groups are stratified based on K-means clustering. Here, the optimal number of clusters (n = 3) was selected automatically by employing an Akaike information criterion, and again separates patients into short, intermediate, and prolonged time on treatment. **D,** summarizes the variable importance of our elastic-net Cox proportional hazard model, after 50 runs and 5-fold cross validation. Here, selection frequency is defined as the proportion of runs in which each gene achieves a coefficient greater than zero, and mean hazard ratio is the mean coefficient across all runs. The PFS of patients receiving first-line CDK4/6i and ET, harboring tumors with pre-treatment functional alterations in TP53 **(E)** and MDM2 **(F)**

To further elucidate the clinical impact of each gene in the broader context of the clinicogenomic feature space, we implemented an elastic net Cox regression on binary pathogenic variant status of each gene as well as select clinical features. Our model identified a “longer response” group (n = 98, 21.9%) from patients with a median progression free survival (PFS) of 35.1 months, compared with an “intermediate” (n = 216, 48.3%, median PFS = 13.8 months) and “short response” group (n = 133, 29.8%, median PFS = 7.81 months) (Fig. 1C). *TP53* and *MDM2* harbored the most important somatic alterations to stratify between these groups, obtaining variable selection frequencies of 1.0 and 0.92 and mean hazard ratios (HR) of 2.02 and 1.39, respectively (Fig. 1D and Fig. S1). Germline pathogenic variants in *BRCA2* were also demonstrated to be an important stratification factor (variable selection frequency 0.96, mean HR of 1.81).

Of these 447 patients, 129 (28.8%) had pre-CDK4/6i loss-of-function variants in *TP53*, corresponding to short PFS (median PFS of 8.7 months, 95% confidence interval [CI]: 6.6 – 10.3 months; HR: 2.13, 95% CI: 1.69 to 2.69; p = 1.99 e - 10; Fig. 1E). Similarly, *MDM2* amplifications were present in 30 patients (6.7%), also corresponding to short PFS (median PFS of 9.1 months, 95% CI: 4.8 – 16.2 months, HR: 1.69, 95% CI: 1.14 to 2.51; p =0.009; Fig. 1F). Clinical analysis suggested that *TP53* loss-of-function variant and *MDM2* amplification affect the long-term response to CDK4/6i; interestingly, both genes are directly involved in regulation of the p53 protein.

### p53 loss or mutation promotes long-term cell outgrowth

The stability of the p53 protein is regulated by MDM2, and overexpressed MDM2 represents a common mechanism of cancer cell inactivation of p53 (*29*). Given that the *TP53* and *MDM2* gene alterations promote inactivation of p53 function, we sought to understand whether and how p53 loss alters CDK4/6i response using isogenic breast cancer cell lines. As the direct target of CDK4/6 kinase is Rb1, we assessed the effect of CDK4/6i treatment on levels of phosphorylated Rb1 (P-RB) in a CDK4/6i sensitive, *TP53*-WT model (MCF7 parental), compared to those lacking p53 (p53KO) or harboring low level of p53 (MDM2 overexpressing) (Fig. 2A and Fig. S1A). We found that 24 hours of abemaciclib (100nM) treatment potently suppressed both P-RB S780 and the expression level of downstream effector E2F in WT, p53KO and MDM2OE cells. However, this effect was not observed in cells intrinsically resistant to CDK4/6i due to high CDK6 expression by inactivation of FAT1 (FAT1KO) (*20*). Accordingly, MCF7 parental, p53KO, and MDM2OE cells all accumulated in G1 (>80%) while FAT1-loss cells did not (Fig. 2B). Further reflecting these initial effects of treatment, the drug concentration (IC50) required to inhibit proliferation was nearly identical for MCF7 parental (9.5nM), p53KO (13.9nM) and MDM2OE (14.9nM) cells (Fig 2C). Similarly, in the CDK4/6i sensitive cell line HCC-1500, p53KO cells showed loss of Rb phosphorylation and G1 arrest comparable to those of parental cells (Fig. S1C and S1D). Previous work has established that HR+ breast cancer cells blocked in G1 can remain effectively inhibited by CDK4/6i for weeks (*20*). To investigate whether mutant *TP53* modulates this effect, we subjected parental and p53 loss cells to prolonged drug treatment for over 5 weeks. Whereas the growth of parental cells remained fully inhibited at 5 weeks, both p53KO and MDM2OE cells resumed proliferation between 4 and 5 weeks, revealing a potential avenue for drug resistance (Fig. 2D, Fig. S1B and Fig. S1E). Similarly, growth of p53KO xenograft tumors was unaffected by CDK4/6i treatment after 5 weeks while WT tumors remained growth suppressed (Fig. 2E). To further ascertain the sufficiency of *TP53* alterations in mediating these effects, we re-introduced wild type or commonly observed *TP53* loss-of-function mutations into p53KO cells and investigated their response to CDK4/6i. Similar to parental MCF-7 and p53 KO models, cells expressing WT, R273H or R280K *TP53* all incurred decreases in P-RB and E2F1 levels (Fig. 2F) and entered G1 cell cycle arrest after 24 hours of CDK4/6i treatment (Fig. 2G). The 5-day IC50 values were nearly equivalent across these three cell lines (WT = 29nM, R273H = 17.3nM, R280K = 33.9nM) (Fig. S2F). However, upon long-term CDK4/6i treatment, the growth of cells with WT *TP53* remained completely inhibited, whereas those with R273H or R280K *TP53* mutants led to outgrowth phenotype (Fig. 2H). Finally, to further assess these phenotypes in a patient derived model, we established breast cancer organoids from patient derived tumor with HR+/HER2- and WT-*TP53* which were sensitive to ribociclib (Fig. S2G and S2H). Compared to non-targeted sgRNA organoids (NTsgRNA), p53 knockout organoids (p53sgRNA) were resistant to continuous abemaciclib treatment (Fig. 2I and Fig. S2I). These findings demonstrate that intact *TP53* is necessary for long-term growth suppression by CDK4/6i in HR+ breast cancer.

**Figure 2.**
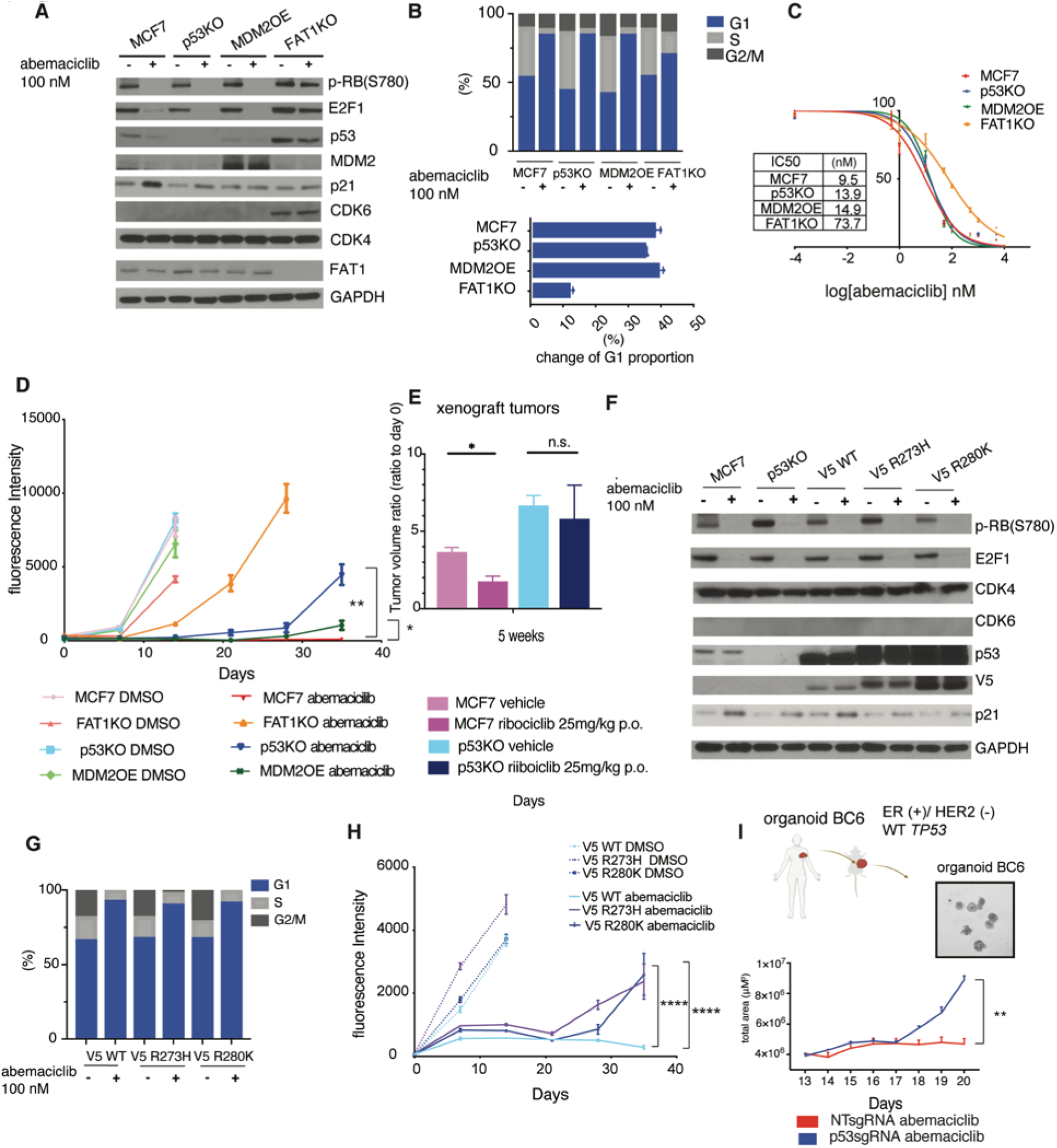
p53 loss promotes long-term cell outgrowth. **A,** Immunoblotting of indicated proteins in parental MCF7 parental (MCF7) cells, cells with knockout of p53 (p53KO), overexpression of MDM2 (MDM 2OE) and knockout of FAT1 (FAT1KO) with 100 nM abemaciclib or DMSO for 24 h. **B,** Cell cycle distribution was measured after 24 h treatment of 100 nM abemaciclib (top). Changes of the proportion of cells in G1 phase (bottom). Data are means ± s.d. of 3 biological replicates. **C,** IC50s of abemaciclib were recorded at day4. **D,** Cell viability assay of parental MCF7, p53KO, MDM2 OE and FAT1KO with 50 nM abemaciclib for indicated days. Data are means of 4 replicates ± SEM. Two-way ANOVA, Tukey’s multiple comparisons test. **E**, Relative tumor volume at 5 weeks compared to tumor volume on day 0 treated with vehicle or ribociclib (25mg/kg) in MCF7 parental and p53KO cells. Data are means of 5 replicates ± SEM. student’s t-test. **F,** Overexpression of V5-tagged wild type p53 (V5 WT), R273H (V5 R273H) and R280K mutants (V5 R280K) in p53 knockout cells. Immunoblotting of indicated proteins with 100 nM abemaciclib or DMSO for 24 h. **G,** Cell cycle distribution was measured after 24h 100 nM abemaciclib treatment. Data are means of 3 replicates. **H**, Cell viability of V5-WT, V5-R273H and V5-R280K cells with 50 nM abemaciclib or DMSO for indicated days. Data are means of 4 replicates ± SEM. Two-way ANOVA, Tukey’s multiple comparisons test. **I,** Organoids BC6 were originally derived from patient tumor with ER+/HER- and WT-TP53. NTsgRNA or p53sgRNA organoids BC6 were treated with 100 nM abemaciclib for 20 days. The area of organoids was measured on indicated days. Data are means of 2 biological replicates ± s.d.. student’s t-test.

### p53 loss enables cell cycle reentry and prevent geroconversion

Akin to non-transformed breast epithelial cells in prolonged G1 arrest, HR+ breast cancer cells have a limited capacity to re-enter the cell cycle and can remain in the arrested state even upon drug withdrawal, where they are subject to the execution of a senescence program that remodels the chromatin and gene expression programs (*5, 15, 30*). We assessed if any of these ‘downstream’ phenotypes might be absent and thus contribute to the outgrowth of p53KO and MDM2OE cells upon CDK4/6i treatment. We examined the effect of drug withdrawal by culturing cells in 50nM abemaciclib for 7 days followed by drug washout and cultured in fresh media. Whereas parental cells showed continuous growth inhibition after drug washout, the p53KO and MDM2OE cultures resumed proliferation and were proficient in colony formation assays compared to MCF7 cells (Fig. 3A and 3B). Similarly, *TP53* knockout and mutant cells retained the potential to generate colonies after drug washout (Fig. S2A). To further ascertain the timing of this effect, we performed FUCCI time-lapse imaging to quantify cell cycle reentry over time following washout of CDK4/6i (*31*). Whereas parental cells showed persistent lack of cells entering S phase over a 75-hour time course, p53KO cells began accumulating in S phase as early as 52 hours after drug washout (Fig. 3C). In addition, we examined the cells for multiple key features of senescence, including expression of senescence-associated-beta-galactactosidase (SA-β-Gal) (Fig. 3D, 3E, 3F and 3K)(*32*), accumulation of senescence associated heterochromatin foci (SAHF) (Fig. 3G, 3H, and Fig. S3B) (*33*), and expression of senescence associated secretory phenotype (SASP) related genes (Fig. 3I, 3J and S2K) (*34*). After 6 days of abemaciclib treatment, ∼80% of parental MCF7 cells were positive for SA-β-gal, as compared to 6%, 20%, and 4% in p53KO, MDM2OE or FAT1KO cells, respectively (Fig. 3D). Induction of WT but not mutant *TP53* restored SA-β-gal activity to levels observed in MCF7 parental cells (Fig. 3E). These findings were recapitulated in HCC-1500 parental and p53KO cells (Fig. S2C and S2D). To further verify the effect of MDM2 on senescence, we utilized pharmacologic inhibition of MDM2 by Nutlin-3a in combination with CDK4/6i and found that this promoted increases in p53 expression and SA-β-Gal activity and decreases on long-term cell proliferation (Fig. S2E, S2F and S2G). Moreover, after 14 days of abemaciclib treatment, SA-β-Gal expression was potently induced in NTsgRNA organoids but not p53sgRNA organoids (Fig. 3F). We also assessed facultative heterochromatin by measuring HP1γ foci positive cells by immunofluorescence and found accumulation in CDK4/6i treated parental cells but not in p53KO, MDM2OE or FAT1-KO cells (Fig. 3G). HP1 γ foci were induced by re-introduction of WT *TP53* (Fig.3H, Fig. S2B). Finally, we sought to identify the SASP genes induced in HR+ breast cancer cells undergoing senescence in response to CDK4/6i and determine if p53KO suppressed their induction. Given the cell type specificity of SASP, we developed a custom nanoString nCounter gene expression panel with 271 genes to identify SASP genes specifically in HR+ breast cancer (Fig. S2H). Comparisons of senescent-proficient (MCF7 parental) versus senescent-deficient (MCF7-FAT1-KO; MCF7-CDK6 high) cells nominated IGFBP5, CAV1, and GEM as strongly associated with senescence *in vitro* and *in vivo* (Fig S2I, S2J and S2K) (*35*). In comparing MCF7 parental and p53KO cells treated with CDK4/6i for 21 days, we identified robust increases in CAV1, IGFBP5 and GEM in parental cells but not in p53KO cells (Fig. 3I). *In vivo* model, SASP genes and SA-β-Gal staining were not increased in p53KO xenograft tumors compared to MCF7 parental tumors (Fig. 3J and 3K).

**Figure 3.**
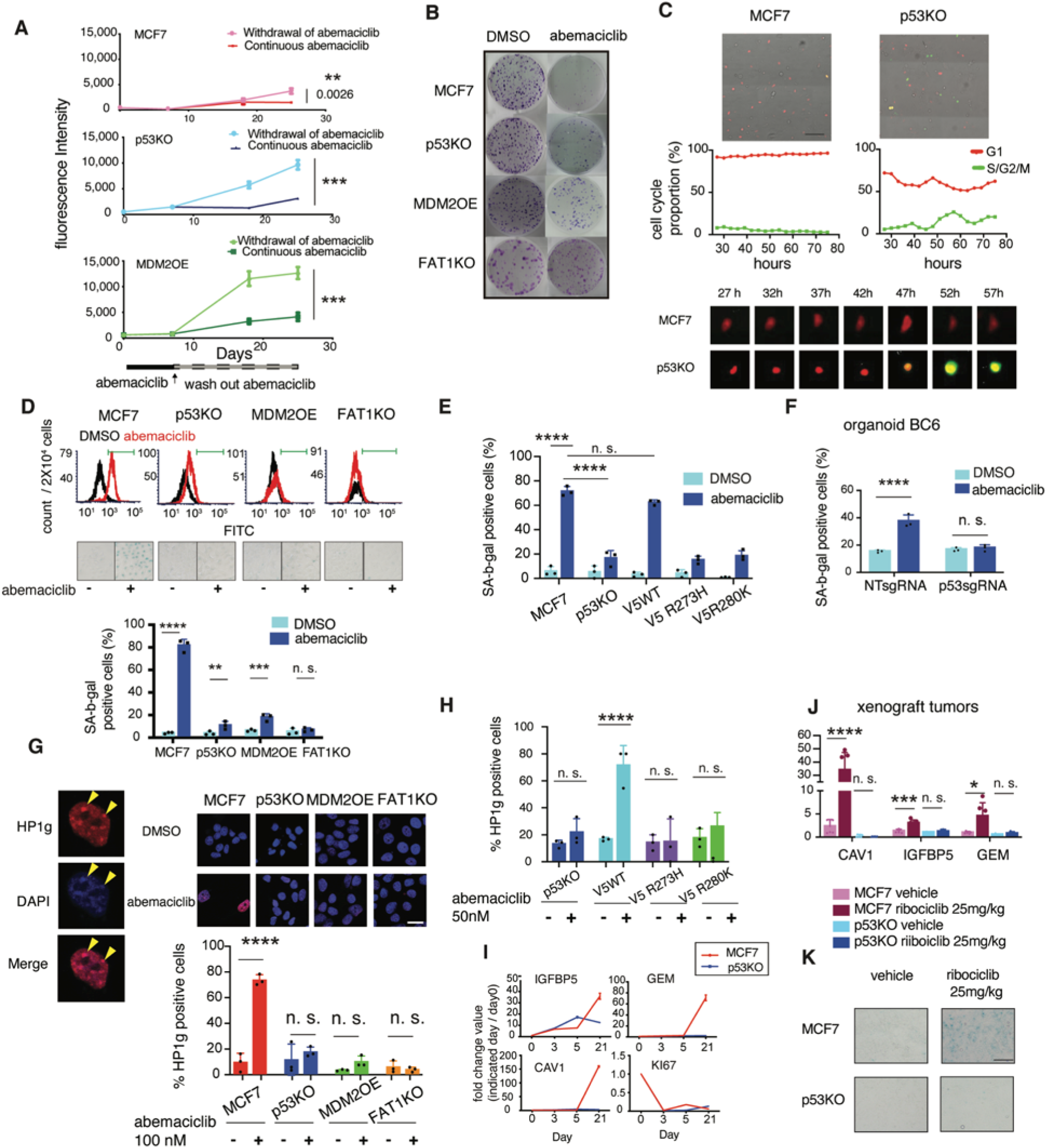
p53 loss enables cells re-enter cell cycle and prevent geroconcersion. **A,** MCF7 parental, p53KO and MDM2 OE cells were treated with 50 nM abemaciclib for 7 days. Cell viability is measured with continuous drug treatment (red, blue and green) or after withdrawal of drug treatment (pink, light blue and light green). Data are means of 4 biological replicates ± SEM. Linear regression analysis was performed. **B,** Each cell line was treated with DMSO or 50 nM abemaciclib for 10 days, and reseeded without drug at 500 cells and cultured for 18 days. **C,** Parental MCF7 and p53KO cell were treated with 100 nM abemaciclib for 6 days. 27h after withdrawal of abemaciclib and simultaneous addition of FUCCI Cell Cycle Sensor, cells were subjected to time-lapse fluorescence microscopy. FUCCI sensors mark cells in G1 phase as red, while cells in S/G2/M are labeled as green. Representative image of each cell lines at 27h after drug withdrawal, bar 100μM (top). Quantification of G1 phase proportion. Data were analyzed by IncuCyte starting after 27h after withdrawal of drug (middle). Representative imaging of MCF7 or 53KO cell starting from G1 phase (red) to G2/M phase (green) every 5 hours (bottom). **D**, Histogram shows counts of SA-β-gal activity positive cells under abemaciclib treatment for 6 days (top). Representative image of SA-β-gal staining in each cell line (middle). Bars represent the percentage of SA-β-Gal positive cells (bottom). Data are means of 3 biological replicates ± s.d.. Two-way ANOVA, Sidak’s multiple comparisons test. **E**, Bars show the percentage of SA-β-Gal positive cells on day 5. Data are means of biological 3 replicates ± SEM. Two-way ANOVA, Sidak’s multiple comparisons test. **F,** Bars represent the percentage of SA-β-Gal positive cells of NTsgRNA or p53sgRNA organoids BC6 after 16 days of DMSO or abemaciclib. Data are means of 3 biological replicates ± s.d.. Two-way ANOVA, Sidak’s multiple comparisons test. **G,** Each cell line was treated with 50 nM abemaciclib for 8 days and the colocalization of HP1γ foci and DAPI was determined by immunofluorescence (yellow arrows). Representative images are shown. Scale bar, 25μM (top). The percentage of HP1γ foci - positive cells was quantified in three different fields. Data are represented as means ± s.d.. Two-way ANOVA, Sidak’s multiple comparisons test (bottom). **H,** V5 WT, V5 R273H, V5 R280K and p53KO cells were treated with 50 nM abemaciclib for 8 days. The percentage of HP1γ foci - positive cells were quantified in three different fields. Data are represented as means ± s.d.. Two-way ANOVA, Sidak’s multiple comparisons test. **I**. MCF7 parental and p53KO cells were treated with 50 nM abemaciclib for indicated days, and RNA-seq was performed. Fold changes (expression on day 3, 5 and 21 /expression on day 0) of IGFBP5, GEM, CAV1 and Ki67 were summarized. (n=2). **J**, qPCR of SASP expressions in MCF7 or p53KO xenograft tumors with vehicle or 25 mg/kg ribociclib treatment after 5 weeks. Relative mRNA expression of indicated genes normalized to PSMC4. The ratios are represented as means ± s.d. from two tumors. n = 6. Student t-test. **K,** Representative images of SA-β-Gal staining of xenograft tumors in J. Bar shows 100μM.

Taken together, these data point to a halt in progression from G1 arrest to senescence in p53 loss cells treated with CDK4/6i, facilitating their re-entry into the cell cycle.

### Loss of p53 suppresses DREAM complex assembly

To identify genes related to the effects of p53 loss upon breast cancer response to CDK4/6i, we analyzed drug-treated parental and p53KO cells over time by RNA-sequencing. Among the most notable gene sets distinct between the models were E2F and DREAM complex targets (Fig. 4A). The DREAM complex is comprised of p130/p107, E2F4, DP1, and a stable core complex of MuvB-like proteins and serves to repress cell cycle dependent gene expression during quiescence (*36, 37*). The DREAM complex can thus maintain cells in quiescence after initial G1 block (*38*). We examined the effect of p53KO upon expression of DREAM complex components after CDK4/6i treatment and found p130 phosphorylation to be discordant between parental and p53KO cells (Fig. 4B). Phosphorylation of p130 impairs its interaction with E2F4, thereby promoting E2F4-mediated transcription and cell cycle reentry (*39, 40*). After 48 hours of abemaciclib treatment, the association between E2F4 and p130 was induced in parental, but not p53KO cells (Fig. 4C). Given the role of p53 in regulating p21 expression and the potential for p21 to suppress p130 phosphorylation (*41, 42*), we engineered a doxycycline-inducible p21 into parental MCF7 and p53KO cells (Fig. 4D). The phosphorylation of p130 persisted in p53KO cells treated with abemaciclib, but was reduced upon induction of p21, resembling the pattern of parental cells. In addition, when stably overexpressing p21, p53KO cells treated with CDK4/6i demonstrated sustained suppression of proliferation and exhibited hallmarks of senescence (Fig. 4E, 4F, and 4G). Time-lapse imaging of p21-expressing cells demonstrates that p21 was sufficient to block cell cycle reentry of p53KO cells treated with CDK4/6i (Fig. 4H). Collectively, these data demonstrate that intact p53 is necessary for DREAM complex assembly in response to CDK4/6i, and that p53 loss can allow cells to reenter the cell cycle and thereby contribute to drug resistance.

**Figure 4.**
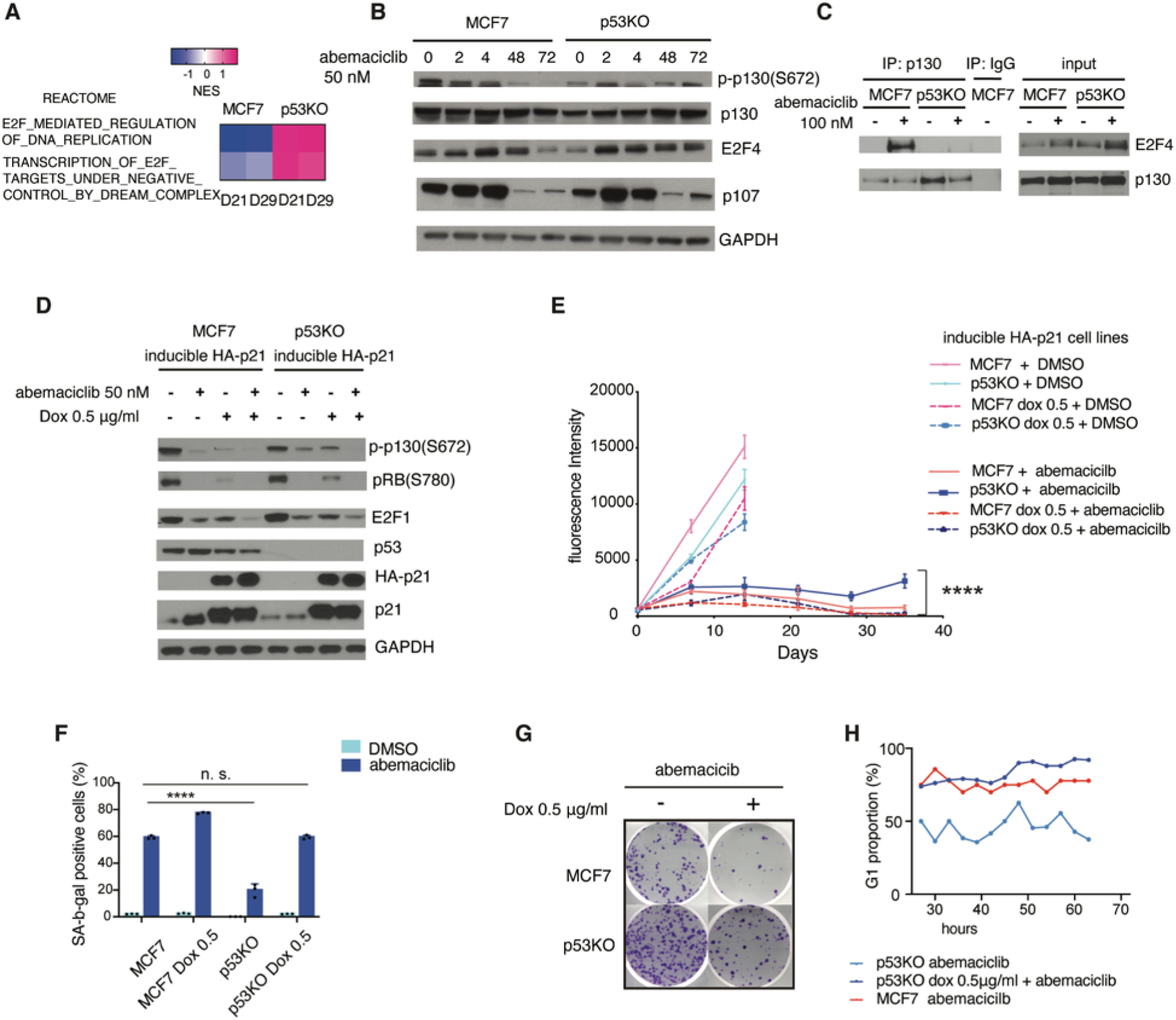
Loss of p53 suppresses DREAM complex assembly. **A,** Heatmap indicating normalized enrichment score (NES) of REACTOME gene sets. NES was analyzed between on day 21 or 29 compared with that on day3. **B,** Immunoblotting of DREAM complex components with 50 nM abemaciclib for indicated hours. **C,** MCF7 and p53KO cells were treated with 100 nM abemaciclib for 48h. Lysates were immunoprecipitated with p130 and IgG antibodies. **D,** MCF7 cells and p53KO transduced with a doxycycline (dox)-inducible HA-tagged p21. The cells were treated with 50 nM abemaciclib or DMSO, ± dox 0.5 μg /ml for 48h. Western blot results with indicated antibodies. **E,** Cell viability of MCF7 parental and p53KO cells with dox-HA-p21, treated with 50 nM abemaciclib or DMSO, ± dox for indicated days. Data are means of 4 biological replicates ± SEM. Two-way ANOVA, Tukey’s test. **F,** SA-β-Gal activity were analyzed by flow cytometry. Bars show the percentage of SA-β-Gal positive cells. Data are means of 3 biological replicates ± s.d.. Two-way ANOVA, Sidak’s test. **G,** Cells were treated with 50 nM abemaciclib ± dox for 11 days, and reseeded after drug washout. **H,** Quantification of G1 proportion in MCF7 without dox or p53KO ± dox. Data were analyzed by IncuCyte starting 27h after withdrawal of drug.

### Combined inhibition of CDK2 and CDK4/6 enables long-term growth suppression

As CDK4/6i leads to down-regulation of CDK2 function through p21 redistribution from CDK4 to CDK2, and p130 can be phosphorylated by CDK2 (*41, 43, 44*), we hypothesized that combined CDK2 and CDK4/6 inhibition might suppress p130 phosphorylation in p53KO cells. We utilized inhibitors to target CDK2 together with CDK4/6 (ebvaciclib alone – CDK2/4/6 inhibitor; PF-07104091 and SNS032 – CDK2 inhibitors; combined with abemaciclib) in MCF7 parental and p53KO cells (*45, 46*). Whereas CDK4/6 inhibition alone was sufficient to inhibit both RB and p130 phosphorylation in parental cells, p53KO cells required the addition of a CDK2 inhibitor to downregulate p130 (Fig. 5A and 5D). Combined inhibition of CDK2 with CDK4/6 enabled long-term growth suppression of p53KO cells (Fig. 5B and 5E) and was associated with potent induction of senescence phenotypes such as SA-β-Gal expression (Fig. 5C and 5F). To examine the combination effect in a patient derived breast cancer model, we established a primary breast cancer cell line from an ER+/HER2- breast cancer that had a loss of function *TP53* mutant, H179R. Combined CDK4/6 and CDK2 inhibition suppressed the phosphorylation of both RB and p130, as well as long-term growth in this *TP53* mutant primary cell line (Fig. 5G and 5H). Moreover, CDK2/4/6 inhibition more potently induced SA-β-Gal expression than abemaciclib did (Fig. 5I). Together, these data reveal that p53KO cells poorly undergo geroconversion in response to CDK4/6i and that this response may be significantly embellished through addition of a CDK2 inhibitor.

**Figure 5.**
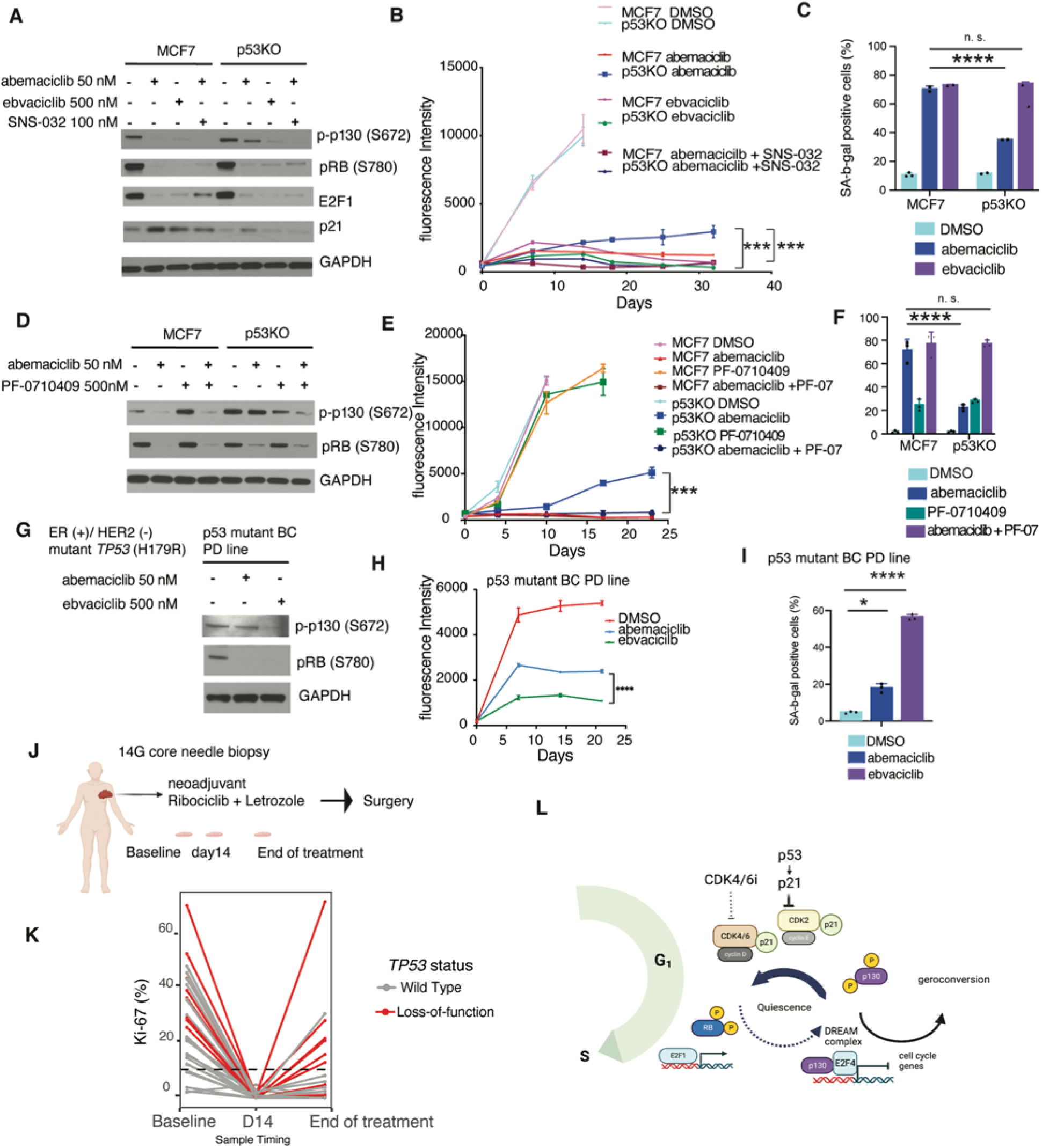
Combined inhibition of CDK2 and CDK4/6 enables long-term growth suppression. **A,** Immunoblotting of MCF7 and p53KO treated with 50 nM abemaciclib, 500 nM ebvaciclib (CDK2/4/6i) or 50 nM abemaciclib±100 nM SNS-032 (CDK2i) for 48h. **B,** Cell viability of MCF7 parental and p53KO cells treated with 500 nM ebvaciclib or 50 nM abemaciclib with or without 50 nM SNS-032 for indicated days. Data are means of 4 biological replicates ± SEM. One-way ANOVA followed by Tukey’s test. **C,** SA-β-Gal activity were analyzed by flow cytometry. MCF7 and 53KO cells treated with 50 nM abemaciclib or 500 nM ebvaciclib for 5 days. Data are means of 3 biological replicates ± s.d.. Two-way ANOVA, Tukey’s test. **D,** Immunoblotting of MCF7 and p53KO treated with DMSO, 50 nM abemaciclib, 500 nM PF-0710409 (PF-07) or combination for 72h. **E,** Cell viability of MCF7 parental and p53KO cells treated with DMSO, 500 nM PF-07, 50 nM abemaciclib or combination for indicated days. Data are means of 4 biological replicates ± SEM. One-way ANOVA followed by Tukey’s test. **F,** SA-β-Gal activity were analyzed by flow cytometry. MCF7 and 53KO cells treated for 6 days. Data are means of 3 biological replicates ± s.d.. Two-way ANOVA, Tukey’s test. **G,** p53 mutant BC primary cell line (*TP53* H179R) were treated with 50 nM abemaciclib and 500 nM ebvaciclib for 48h. Western blot results with indicated antibodies. **H,** Cell viability of p53 mutant BC PD line treated with 500 nM ebvaciclib or 50 nM abemaciclib. Data are means of 4 biological replicates ± SEM. One-way ANOVA followed by Tukey’s test. **I,** SA-β-Gal activity in p53 mutant BC PD line cells were analyzed by flow cytometry. Bars are showing the percentage of SA-β-Gal positive cells. Data are means of 3 biological replicates ± s.d.. Two-way ANOVA, Sidak’s test. **J,** Schematic diagram of sample collection in FELINE trial. Core needle biopsy was performed over the course of treatment: screening (day 0), cycle 1 day 14 (day 14) and end of trial (day 180) using a 14-gauge needle. **K,** Ki-67 trend is depicted for all patients in the FELINE trial for whom Ki-67 was evaluable at all time points and sequencing of the day 0 tumor was performed. The rates of Ki-67 > 10% at surgery are compared by pre-treatment *TP53* status utilizing Fisher’s Exact (two-sided) test. L, Model for loss of p53 resistant mechanism. In the presence of p53, CDK4/6i inhibited phosphorylation of RB and p130. The cells were arrested in quiescence through DREAM complex, leading to irreversible cell cycle arrest. Loss-of-function p53 and insufficient p21 enabled CDK2 to retain phosphorylated p130, leading to cell cycle reentry and the escape from quiescence.

### p53 loss is associated with cell cycle reentry in patients treated with neoadjuvant CDK4/6i

Finally, to assess the relevance of these results to human breast cancer patients, we examined the impact of *TP53* status upon patients treated with neoadjuvant ribociclib on the FELINE clinical trial (NCT0271273). In this trial, two cohorts of patients with early stage ER+ breast cancer were treated with combinations of the CDK4/6i ribociclib with aromatase inhibition and evaluated at two weeks and 180 days on therapy for changes in Ki67 (Fig. 5J) (*47*). Pretreatment and on-treatment samples from 33 patients were subjected to whole genome sequencing. Change in Ki-67 (dichotomized at 10%) on neoadjuvant endocrine therapy has been associated with improved time to recurrence in prior neoadjuvant endocrine trials and thus utilized in this cohort (*48*) (*49*). Among 33 evaluable patients, 10 (30.3%) harbored a pre-treatment *TP53* loss of function variant. Of these 10 cases, 6 (60%) did not achieve a low (< 10%) Ki-67 upon surgical resection as compared to *TP53* wildtype tumors (n=23), only one (3.2%) of which did not achieve a low Ki-67 (OR 28.1, 95% CI 2.51 – 1564, p = 0.0012) (Fig. 5K). These data provide further support that p53 loss prevents a durable cell cycle arrest in ER+ breast cancer thereby limiting the full therapeutic potential for this approach.

## Discussion

In this report, we investigated the underlying basis for long-term response to CDK4/6 inhibition and identify defective p53 function to be a highly prevalent determinant preventing long-term disease control. While identifying the clinical scenario of high-risk or metastatic ER+ breast cancer defines the group of patients that can benefit from CDK4/6 inhibitors, it fails to establish which patients have a high likelihood of long-term clinical benefit owing to the wide variance in duration of response observed (*50*). Our data demonstrate that while the proximal effect of the drug at inhibiting its target and inducing G1 arrest is more widely achieved, the ability of the drug to maintain cancer cells in quiescence via the DREAM complex and initiation a senesence program separates the cancers likely or not to have long-term disease control. Critically, we identify a highly prevelant genetic alteration, p53 loss, to mediate these differences and to be potentially targetable via an emergent class of treatments, selective CDK2 inhibitors.

To uncover the clinically-relevant ‘requirements’ for long-term disease control on CDK4/6i therapy, we surveyed for genetic alterations specifically absent in patients who have been on prolonged therapy with CDK4/6i. We and others have previously identified a number of genetic alterations that mediate *de novo* resistance to CDK4/6i therapy (*20, 35, 51*), but these mainly constitute uncommon alterations that promote immediate lack of drug response. By contrast, the majority of patients who go on CDK4/6i therapy are on therapy for >6 months and we thusly focused our efforts at identifying genetic alterations that separate an ‘intermediate’ response from a more durable response. Using these criteria and a database of 447 patients treated with first-line CDK4/6i, we identified pre-existing loss-of-function variants in *TP53* as under-represented in patients who experience prolonged response. *TP53* is one of the most commonly mutated genes in breast cancer and plays protean roles in tumor maintenance and progression, regulating DNA damage repair, cell cycle, chromatin architecture, cellular metabolism, cell invasion, immune responses, and cell death (*21, 52, 53*). Thus, the ways in which mutant *TP53* might be relevant to drug response might be multifactorial. However, our findings on CDK4/6i response point to the specific function of p53 in regulating the DREAM complex in mediating these differences in outcomes. In further support of a central role for p53 in this effect, alterations in genes known to interact with *TP53, MDM2* and *PPM1D,* were also enriched in patients who did not achieve a long-term response (*54, 55*). These findings are consistent with the prior observations of a prognostic role for *TP53* mutations in HR+ breast cancer (*56*) and can immediately facilitate efforts at risk stratifying patients with high-risk vs low-risk advanced disease through enhanced surveillance and novel drug combination strategies for patients with *TP53* loss-of-function mutant disease.

In order to discover the biologic mechanisms underpinning therapy failure in p53 mutant tumors, we analyzed both the short-term and long-term response of isogenic models to CDK4/6i. While CDK4/6i uniformly led to reduced Rb1 phosphorylation and G1 arrest in p53 proficient and deficient cells, we found that over the long-term, p53 deficient cells ultimately developed resistance and resumed proliferation. Strikingly, p53 deficient cells failed to initiate the changes that lead to phenotypes of cancer cell senescence, changes that were uniformly seen in the p53 proficient models. Mirroring these results from breast cancer models, surveying breast cancer patients treated with ribociclib on the FELINE clinical trial revealed that resumption of the cell cycle after initial G1 arrest was commonly observed in p53 mutant cases but not in p53 wild type cases. The specific contribution of the senescence program to mediating the efficacy of the CDK4/6i is uncertain as there may be both tumor suppressive and tumor promoting factors that can be elicited in different contexts (*19, 57*). Nevertheless, failure-to-induce senescence in this scenario was reflective of a capacity of p53 deficient cells to re-enter the cell cycle and thereby reinitiate proliferation. Moreover, we found nearly identical results in modeling MDM2 overexpression and its resulting effect of lowering p53 expression levels. As there may be several genetic changes in addition to p53 loss that result in the same impairment of long-term growth suppression, the SASP phenotype could serve as useful phenotypic biomarker for detecting cancers highly responsive to G1 checkpoint directed therapies.

The failure of p53KO cells to elaborate any of the senescence changes suggested that the defect in this context was upstream of the epigenetic and transcriptional re-programming that mediate these phenotypes. We examined the gene expression changes in response to CDK4/6i in these cells and identified changes in E2F and DREAM complex targets as differential between the p53 proficient and deficient models. Based on the seminal work identifying the function of the DREAM complex in maintaining cellular quiescence and blocking cell cycle re-entry (*36*), we analyzed the assembly of this complex and identified p130 as differentially phosphorylated in p53 KO and WT drug-treated cells. The lower levels of p21 expressed in p53 KO cells provided a probable candidate for residual p130 phosphorylation in the CDK2 kinase. Either reestablishment of p21 expression or direct inhibition of CDK2 was capable of facilitating p130 de-phosphorylation, blocking cell cycle reentry, and enabling durable inhibition of cell proliferation (Fig. 5L). These findings serve to highlight the critical function of the DREAM complex in mediating the antitumor effects of G1 directed anticancer therapeutics and point to a potentially important indication for several novel and highly selective CDK2 inhibitors now entering the clinic (*45, 58*). After determining the safety and therapeutic index of these drugs, our data imply that the p53 deficient set of tumors (a highly prevalent subset) may represent a major indication, serving to distinguish them from the current generation of CDK4/6i and potentially extending the benefit of G1-directed therapies to a significantly wider population of patients.

## Materials and Methods

### Cell lines

MCF7, HCC1500 cell lines were obtained from the American Type Culture Collection. HEK293T was a gift from Ping Chi’s lab. MCF7 cells were maintained in DMEM/F12 medium. HCC1500 cells were maintained in RPMI medium. All media were supplemented with 10% FBS, 2 mmol/L L-glutamine, 20 U/mL penicillin, and 20 μg/mL streptomycin. All cell lines were authenticated by short tandem repeat genotyping and tested negative for Mycoplasma contamination. All cell lines were cultured and collected within 10 passages.

### Drugs and Reagents

Abemaciclib (LY2835219), palbociclib (PD-0332991), Nutlin-3a (S8059), PF-07104091(S9878) and SNS-032 (BMS-387032) were obtained from Selleck Chemicals. Ebvaciclib (PF-06873600) was obtained from AdooQ BioScience. We obtained ribociclib (LEE011) from Novartis. These drugs were dissolved in dimethylsulfoxide.

### Antibodies

Phospho-RB1 (Ser780; #8180), phospho-RB1 (Ser807/811; #8516), RB1 (#9309), cyclin D1 (#2978), CDK6 (#3136), CDK4(#12790), CDK2 (#2546), E2F1 (#3742), cyclin A2 (#4656), cyclin E2 (#4132), GAPDH (#5174), Rbl1 (#89798), Rbl2 (#13610), E2F4 (#40291), p53 (#9282), p21 (#2947), HA-Tag (#3724), V5-Tag (#13202) and MDM2 (#86934) antibodies were purchased from Cell Signaling Technology. Phospho-p130 (S672) (#ab76255) antibody were purchased from Abcam. P130 (A-10) and normal mouse IgG (sc-2025) were purchased for immunoprecipitation from Santa Cruz Biotechnology. MDM2 (OP46) was purchased from EMD Millipore.

### Breast cancer organoid establishment

PDX tissue was washed with cold PBS containing antibiotics upon arrival and cut into 1-3mm3 pieces. Small pieces were digested using tumor dissociation kit (Miltenyi Biotec, 130-095-929) on gentleMACS Octo Dissociator at 37C for 1h. Digested tissue suspension was spun down at 600 rcf for 5 mins. Aspirate off the supernatant before adding 10 ml TrypLE (Gibco, 12563011) and incubate at 37 for 10 mins on thermomixer. The suspension was strained over a 100 um filter. Add 10 ml Advanced DMEM/F12 containing 1x Glutamax, 10mMHEPES, and antibiotics to rinse off the tube. Repeat this step if necessary. Spin down at 600 rcf for 5 mins. Aspirate off the supernatant before adding 1 ml of ACK lysis buffer (Gibco, A1049201) and incubate for 5 mins at room temperature. Add PBS to stop the reaction. Spin down at 600 rcf for 5 mins. The pellet was resuspended in 1 ml of cold Matrigel (Corning, 356231). Add 40 ul of suspended samples to each well of prewarmed 24well plates (Greiner, M9312). Solidified in 37C after 20 mins, 400ml warm of organoid media was added to each well. Media were refreshed every 3 or 4 days and organoid were passaged every 9 days. Organoid culture media contains Advanced DMEM-F12 without phenol red (Gibco), 100× Glutamax (Life teqnologies, 35050-061), 10mM HEPES (Gibco, 15630080), 50x B27 supplement (Gibco, 17504044), 100x Penicillin-Streptomycin(Gibco), 100 μg/mL primocin (Invitorgen, NC9141851), 10% R-Spondin conditioned medium, 10% Noggin conditioned medium, 500 nmol/L A83-01 (Tocris Bioscience, 29-391-0), 1ng/mL hEGF (PeproTech, AF-100-15), 100 ng/mL Hydrocortisone (Sigma Aldrich, H0888), 5ng/mL hFGF2 (PeproTech, AF-100-18B), 5ng/mL hFGF7 (PeproTech, 100-19), 10ng/mL hFGF10 (PeproTech, AF-100-26), 1 nmol/L estradiol, 1 ng/mL Neuregulin 1 (PeproTech, 100-03), 1.25 mmol/L N-acetylcysteine (Sigma Aldrich, A9165), 10mol/L nicotinammide (Sigma Aldrich, N0636), 4μg/mL Heparin (Stemcell technologies) and 5 μmol/L Y-27632 (Enzo Life Science, 50-103-1737).

### Primary breast cancer cell line establishment

PDX tumor for p53 mutant BC PD line was washed with cold PBS containing antibiotics upon arrival and cut into 1-3mm3 pieces. Small pieces were digested using tumor dissociation kit (Miltenyi Biotec, 130-095-929) on gentleMACS Octo Dissociator at 37C for 1h. Digested tissue suspension was spun down at 600 rcf for 5 mins. Aspirate off the supernatant before adding 10 ml TrypLE (Gibco, 12563011) and incubate at 37 for 10 mins on thermomixer. The suspension was strained over a 100 um filter. Add 10 ml Advanced DMEM/F12 containing 1x Glutamax, 10mMHEPES, and antibiotics to rinse off the tube. Repeat this step if necessary. Spin down at 600 rcf for 5 mins. Aspirate off the supernatant before adding 1 ml of ACK lysis buffer (Gibco, A1049201) and incubate for 5 mins at room temperature. Add PBS to stop the reaction. Spin down at 600 rcf for 5 mins. Cells were resuspended medium and seeded.

### SA-β-gal assays

Cells were treated with DMSO, 50 nM abemaciclib or 500 nM evbaciclib for 5 −8 days. Cells were harvested using trypsin/EDTA and then stained with the CellEvent Senescence Green Flow Cytometry Assay Kit (Invitrogen, C10840) according to the manufacturer’s instructions. Briefly, cells were fixed with 2% paraformaldehyde for 15 minutes at room temperature and stained with the CellEvent Senescence Green Probe for 2 hours at 37°C without CO2. After incubation, cells were washed with PBS three times and resuspended in FACS buffer (PBS with 2%FBS) for analysis on BD Biosciences LSR Fortessa using a 488-nm laser and 530-nm/30 filter (BD Biosciences). Data analysis was performed with FCS Express 7 (De Novo Software). SA-β-gal staining was assayed using Senescence β-Galactosidase Staining Kit from Cell Signaling Technologies (#9860) according to manufacturer’s instructions and previously described (*19*). For tumor tissue staining, 10μm thickness slides were prepared from OCT compound frozeh tissue and stained for over night with the kit.

### Cell-cycle Analysis

Cells were treated with DMSO or 100 nM abemaciclib for 24 hours. Cells were detached from the cell culture dish with trypsin/EDTA, and then washed with PBS and fixed in 70% ice-cold EtOH overnight. Prior to staining, EtOH was removed, and cells were washed twice with FACS buffer. Cells were then resuspended in staining buffer containing 1,000 μL FACS buffer with 2 μg/mL propidium iodide (Invitrogen) and 100 μg/mL Rnase A (Invitrogen). Cell-cycle profiles were measured with BD Biosciences LSR Fortessa and analyzed with FCS Express 7.

### Cloning and Plasmids

LentiCRISPRv2 puro backbone was used for generating knockout cell lines. lentiCRISPRv2 puro was a gift from Brett Stringer (Addgene plasmid # 98290). Single-guide RNAs were designed through MIT CRISPR Designer (crispr.mit.edu), and the sequences are as follows: p53-CRISPR, CACCGGAGCGCTGCTCAGATAGCGA; Instructions for using the lentiCRISPRv2 plasmids are as described by the Zhang laboratory (https://media.addgene.org/cms/filer_public/53/09/53091cde-b1ee-47ee-97cf-9b3b05d290f2/lenticrisprv2-and-lentiguide-oligo-cloning-protocol.pdf). Oligos were annealed and ligated with BsmBI-digested lentiviral vector. Then, the ligation system was transformed into Stbl3 bacteria, and plasmids were extracted for sequencing.

For another guide RNA, we obtained sgTP53_3 which was a gift from William Hahn (Addgene plasmid # 78164), and lentiCRISPRv2 hygro which was a gift from Brett Stringer (Addgene plasmid # 98291).

pDONR223_MDM2_WT was a gift from Jesse Boehm & Matthew Meyerson & David Root (Addgene plasmid # 82897) cloned into pLenti PGK Neo DEST (pLenti PGK Neo DEST (w531-1) a gift from Eric Campeau & Paul Kaufman (Addgene plasmid # 19067) by using the Gateway LR Clonase II Enzyme Mix (Invitrogen).

pDONR223_CDKN1A_WT was a gift from Jesse Boehm & William Hahn & David Root (Addgene plasmid # 82201) was cloned into pInducer20 (a gift from Stephen Elledge, Addgene plasmid # 44012) by using the Gateway LR Clonase II Enzyme Mix. As p53 expressing vector, pLenti6/V5-p53_wt p53, pLenti6/V5-p53_R273H and pLenti6/V5-p53_R280K were used. These plasmids were gifts from Bernard Futscher (Addgene plasmid # 22945, # 22934 and # 22933). tFucci(CA)5 was a gift from Atsushi Miyawaki (Addgene plasmid # 153521).

### Lentiviral and retroviral infection and generation of stable cell lines

HEK293T cells were transfected with 4.5µg of lentiviral vector, 4.5 µg of psPAX2 and 1µg of pVSVG with 40µl X-tremeGENE HP (Roche) according to the manufacturer’s protocol. Conditioned medium containing recombinant lentivirus was collected 48hrs after transfection and filtered through non-pyrogenic filters with a pore size of 0.45 μm (Merck Millipore). Samples of these supernatants were applied immediately to target cells together with Polybrene (Sigma-Aldrich) at a final concentration of 8 μg/ml, and supernatants were incubated with cells. After infection, cells were placed in fresh growth medium and cultured as usual. Selection with 2 µg/ml puromycin (Thermo Fisher Scientific) for 7 days, 1 mg/ml G418 (InvivoGen) for 14 days or 200 µg/ml hygromycin (InvivoGen) for 7 days was initiated 48 h after infection for MCF7. Selection with 0.5 µg/ml puromycin was initiated 72h after infection for HCC1500.

### Cell viability assay

For cell lines, cell viability was measured by Resazurin (R&D Systems, Minneapolis, MN, USA) as described previously (*20*). Briefly, 1,500 cells were seeded to 96-well plate allowed to recover overnight. Cells were treated with drugs at day 0. Resazurin was added to the cells 4 hours prior to the measurement at day 3, day 5 and day 7. For long-term growth assay, 300 cells were seeded in 96-well. Fluorescence in the plate was measured using a microplate reader (SpectraMax M5, Molecular Devices). IC50 was calculated by GraphPad Prism 9.0 by using a sigmoidal regression model. For organoids, cell viability was determined over time course with a a microplate reader using CellTiter-Glo® 3D Cell Viability Assay (Promega, G9683) according to manufacturer’s directions. The area of organoids was analyzed using the Incucyte S3 (Sartorius).

### Immunoblotting

Cells were lysed in RIPA buffer (Thermo Scientific, 89901) supplemented with 1x protease and phosphatase inhibitor cocktail (Thermo Scientific, 78445). Protein concentration was determined using the BCA protein assay kit (Thermo Scientific, 23227) and equalized using RIPA buffer. Cell lysates were subjected to 4-12% Bis-Tris gel (Thermo Scientific, WG1402BOX) electrophoresis, transferred to nitrocellulose membranes and blocked with 5% non-fat milk in TBST for 1 hr at room temperature. Membranes were incubated with the indicated antibodies overnight and then with secondary HRP-coupled anti-mouse (CST, #7076) or anti-rabbit antibody (CST, #7074) for 1 hr at room temperature and visualized using Konica Minolta Film Processor with Western Lightning Plus (Perkin Elmer, NEL104001) or Immobilon Crescendo Western HRP substrate (Millipore, WBLUR0100).

### Colony formation assay

MCF7 cells were treated with DMSO or 50 nM abemaciclib for 10 days, and 500 cells were seeded in 6 well plate after drug withdrawal for colony formation assay. HCC1500 cells were treated with DMSO or 100 nM abemaciclib for 10 days and seeded at 50000 cells in 12 well plate. Cells were cultured without drug for 18-21 days. After removal of medium, 100% methanol were added for fixation and cells were incubated for 20 mins. After removal of methanol and the cells were rinsed with H2O and covered with crystal violet staining solution (0.5% crystal violet in 25% methanol) for 5 min. The cells were washed with H2O twice and then dried up overnight.

### Immunoprecipitation

Cells were lysed in IP lysis buffer (Pierce, 87787) supplemented with 1x protease and phosphatase inhibitor cocktail (Thermo Scientific, 78445). The antibody for p130 and mouse IgG were incubated with the Dynabeads Protein G (Invitrogen, 10003D) for 4 hrs at 4°C. Excess antibody is washed away by placing the tube in a DynaMag magnet and removing the supernatant. Lysate and Dynabeads were incubated overnight with end-over-end agitation at 4°C. Beads were separated by magnet and washed with IP lysis buffer 3 times before proteins were eluted by heating in 2xNuPAGE LDS sample buffer (Invitrogen, NP0008) and 2x sample reducing buffer (Invitrogen, NP0009) with IP lysis buffer for 5 minutes at 70 °C. Samples were subjected to downstream western blotting with 8% Bis-Tris gel (Thermo Scientific, NW00080BOX) electrophoresis.

### FUCCI cell cycle time-lapse imaging

MCF7 or p53KO MCF7 cells were treated with 50 nM abemaciclib for 6-10 days, and replated in Lab-Tek™ II Chambered Coverglass (Thermo scientific, 155382PK) without drug. On the same day, Invitrogen™ Premo™ FUCCI Cell Cycle Sensor (Invitrogen, P36238) was added to chamber. After 27 hours, images were taken every hour with a Zeiss Axio Observer Z1 inside a heated 37°C chamber with 5% CO2 for 48 hours and analyzed by ZEN Blue 3.2 software. HA-p21 dox inducible cell line which stably expressed tFucci (CA)5, were seeded in Lab-Tek™ II Chambered Coverglass and treated with 50 nM abemaciclib. After 24 hours, 0.5*µ*g/ml doxycycline was added to chamber to induce p21. After 4 days abemaciclib treatment, medium was refreshed with 0.5*µ*g/ml doxycycline and without abemaciclib. The images were taken every 3 hours for 72 hours.

To quantify the cell cycle phase proportion, MCF7 and p53KO cells were plated in 6cm dish and IncuCyte Cell Cycle Green/Red Lentivirus Reagents (Sarutorius, 4779) and polybrene were added. After 2 µg/ml puromycin selection for one week, the cells with fluorescence were sorted by Aria-3 cell sorter (BD bioscience). Sorted cells were seeded in Corning 96 well plates (Corning, 3595) and treated with 50 nM abemaciclib for 6 days. The medium was refreshed without drug. Images are acquired and analyzed by using a Sartorius IncuCyte S3 with 3 hours imaging intervals. For HA-p21 dox inducible cell lines which were expressing tFucci(CA)5, IncuCyte S3 was used directly to analyze cell cycle analysis.

### Senescence-associated heterochromatin foci (SAHF) assay

Cells were plated on 4-well Millicell EZ SLIDE glass (EMD Millipore, PEZGS0416). After 7-10 days DMSO or drug treatment, cells were and fixed for 10 min with 4% paraformaldehyde dissolved in PBS. After three times wash in PBS, cells were incubatoed in 1% BSA dissolved in PBST for 30 mins. Cells were then incubated with HP1g antibody (anti-HP1*y* antibody, clone 42s2, EMD Millipore, 05-690, 1:5000) at 4°C overnight, prior to washing with PBS and incubation with anti-mouse secondary antibodies (Goat anti-Mouse IgG (H+L) Cross-Adsorbed Secondary Antibody, Alexa Fluo 594,1:500) for 1 hour at room temperature. After further washing, coverslips were mounted onto slides with Mounting Medium with DAPI (abcam, ab104139) covered with Fisgerbrand Premiun Cover Glasses (Fisher Scientific, 12-548-5M). Images were acquired Leica TCS SP5 Upright microscope, and analyzed with the LAS AF software.

### In Vivo Tumor Models

Tumors were obtained from MCF7 parental cells, MCF7-p53KO or MCF7-CDK6 expressing cells xenograft tumors which were previously reported (*20*). Briefly, NOD.Cg-Prkdc <scid> Il2rg<tm1Wjl>/SzJ (NSG) mice were obtained from the Jackson Laboratory (stock no. 005557). Each mouse was injected with MCF7 parental or CDK6-ovexpressing, subcutaneously 1 week after the implantation of estradiol pellets (25 mg). Each mouse was injected MCF7 parental or MCF7-p53KO. 0.18mg b-estradiol pellets 3 days prior to cell implantation. Mice were treated at a 5 days on/2 days off schedule for 25 to 35 days with ribociclib at 25 mg/kg (orally). BC6 PDX tumor was provided by Harikrishna Nakshatri. 0.18mg b-estradiol pellets 3 days prior to tumor implantation. Mice were treated at a 5 days on/2 days off schedule for 25 to 35 days with ribociclib 75 mg/kg (orally). The study was approved by the IACUC review board at MSK.

### Quantitative RT-PCR

Quantitative RT-PCR was performed as described previously (*20*). Briefly, total RNA was extracted using Rneasy Mini Kit (Qiagen, 74106). RNA was reverse-transcribed into cDNA using qScript cDNA SuperMix (Quanta Biosciences, 95048). qPCR reactions were performed with TaqMan PCR Master Mix (Applied Biosystems, 4369016) using a ViiA 7 Real-Time PCR system (Applied Biosystems) along with primers. Samples were run in triplicate, and mRNA levels were normalized to those of RPLP0 or PSMC4 for each reaction. Taqman primers were purchased from Applied Biosystems and included: GEM (Hs00170633_m1), PSMC4 (Hs00197826_m1), RPLP0 (Hs99999902_m1), IGFBP5 (Hs00181213_m1), CDKN1A (Hs00355782_m1), MKI67 (Hs01032443_m1), E2F1 (Hs00153451_m1), CAV1 (Hs00971716_m1), TBP (Hs00427620_m1).

### RNA sequencing

Cells were treated with 50nM abemaciclib for 0h, 1 day, 3 days, 5 days, 21 days and 29 days. Total RNA from these cells was extracted using Rneasy Kit (QiagenTwo independent biological replicates of each RNA sample were submitted for library preparation and RNA sequencing. RNAs were sequenced by Illumina HiSeq 2×150 bp sequencing in Genewiz. Raw sequencing reads were aligned to the human genome (GRCh38.74) using STAR. Raw feature counts were normalized and differential expression analysis using DESeq2. Differential expression rank order was utilized for subsequent Gene Set Enrichment Analysis (GSEA), performed using the fgsea (v1.8.0) package in R (v 4.2.0). Gene sets queried included those from the Reactome Gene Sets available through the Molecular Signatures Database (MsigDB).

### Nanostring nCounter assay

The nCounter platform (NanoString Technologies) was used to analyze RNA samples cell lines. A minimum of 100 ng of total RNA was used to measure the expression of 271 genes of custom nanoString nCounter gene expression panel and 12 housekeeping genes. Expression counts were then normalized using the nSolver 4.0 software (nanoString, Seattle, WA, USA)

### Statistical analysis

Analyses were conducted using GraphPad Prism 9 and Rstudio. The numbers of biologically independent experiments, the number of individual mice or patient numbers, details of statistical tests performed can be found in the respective figure legends or method section. We showed P values *P < 0.05, **P < 0.01, ***P < 0.001, **** P< 0.0001 in figures.

### Human Subjects

A total of 297 patients with metastatic HR+ breast cancer treated with CDK4/6 inhibitors in combination with hormonal therapy the first-line setting underwent prospective clinical genomic profiling between April 2014 and September 2022. This study was approved by the Memorial Sloan Kettering Cancer Center Institutional Review Board (IRB) and all patients provided written informed consent for tumor sequencing and review of patient medical records for detailed demographic, pathologic, and treatment information (NCT01775072). All protein-coding exons and selected intronic and regulatory regions of 341 to 468 cancer-associated genes, all as previously described (*21*) (*22*) (*23*). Somatic mutations, DNA copy-number alterations, and structural rearrangements were identified as previously described and all mutations were manually reviewed. For each gene, oncogenic relevance of specific variants was assessed using the latest versions on the OncoKB knowledgebase (*24*). For each patient with genomic sequencing performed on a tumor sample acquired before the initiation of CDK4/6i, the presence or absence of a putatively gain or loss-of-function variant was noted. For each patient with genomic sequencing performed on a tumor sample acquired before the initiation of CDK4/6i and/or endocrine therapy, the presence or absence of a predicted oncogenic variant in any pretreatment sample was annotated. Genes with predicted oncogenic alterations in < 5% of patients were filtered out of subsequent analyses.

We categorized CDK4/6 inhibitor regimens based on their endocrine therapy backbone to two major categories: 1) CDK4/6 inhibitors plus aromatase inhibitors, including letrozole, exemestane, or anastrozole and 2) CDK4/6 inhibitors plus selective estrogen receptor degraders (SERD) including fulvestrant. We divided time on treatment into tertiles to distinguish between short, intermediate and long-term responders. To isolate the effects between patients who failed to achieve long-term response, we excluded short term responses (first tertile) and patients in the second tertile who stopped therapy for reasons other than disease progression.

For each gene, we iteratively performed Firth penalized logistic regression, with achievement of long-term response as the dependent variable and presence of pathogenic variant in pre-treatment sample as the independent variable. This was performed in the univariate setting, as well as adjusting for endocrine therapy partner. We rejected the null hypotheses with a two-sided *P* = 0.05. Resulting p values were corrected for multiple hypothesis testing with Benjamini and Hochberg method. For select genes achieving significance, we used univariate Cox Proportional hazard models stratified by treatment class (CDK4/6i + AI or CDK4/6i + SERD) to determine the association between genomic alterations in each gene PFS. We tested the proportionality assumption of the Cox regression model through time-dependency analysis of selected genetic alterations (cox.zph function of the R package survival). All analyses were performed in R version 4.1.0 (R Foundation).

### FELINE trial

#### Sample collection

We studied the breast cancer samples from the FELINE clinical trial (ClinicalTrials.gov identifier: <u>NCT02712723</u>), which evaluated the addition of CDK inhibition to endocrine therapy in the neoadjuvant setting. Patients (*n*□=□120) were randomized equally into three arms: (1) endocrine therapy alone (letrozole plus placebo), (2) intermittent high-dose combination therapy (letrozole plus ribociclib (600 mg□d^−1^, 3 weeks on/1 week off)) or (3) continuous lower-dose combination therapy (letrozole plus ribociclib (400 mg□d^−1^)). Patients were treated for six cycles and biopsies were collected at baseline (SC), following treatment initiation at day 14 (CD) and end of treatment at surgery around day 180 (EOT). Immediately after collection, biopsy samples were snap-frozen and embedded in OCT compound. Informed consent was obtained from all patients following protocols approved by the institutional review boards and in accordance with the Declaration of Helsinki. This study was approved by University of Kansas Institutional Review Board (protocol no. CLEE011XUS10T) (*18*).

#### Sample Processing

Representative sections of 135 OCT compound-embedded core tumor biopsies from 65 patients were reviewed by two pathologists with expertise in breast cancer (Qiqi Ye and J.S. Reis-Filho), and the tumor cell content assessed for subsequent tissue microdissection and DNA extraction.

#### Microdissection and DNA extraction

Eight-μm-thick sections from frozen tumor blocks were stained with nuclear fast red and microdissected using a sterile needle under a stereomicroscope (Olympus SZ61) to enrich tumor content, as previously described (*25*) (*26*). Genomic DNA of microdissected tumor and matched normal samples (buffy coat) was extracted using the DNeasy Blood and Tissue Kit (Qiagen) and QIAamp DNA Blood Mini Kit (Qiagen), respectively, according to manufacturers’ instructions.

#### Whole genome sequencing

DNA from baseline (SC), day 14 (CD), end of treatment (EOT) tumor and matched normal samples were subjected to whole-genome sequencing (WGS) at MSK’s Integrated Genomics Operations (IGO) using validated protocols (*27*) (*28*), with a median sequencing coverage depth of 57x (range, 48x-62x). WGS was completed on 52 SC, 48 CD and 35 EOT tumor and matched normal samples from 65 cases.

## Acknowledgments

We thank the research staffs at Core Facilities at MSK: Eric Rosiek, Eric Chan and YevgeniyRomin (Molecular Cytology), Sydney Bowker, Kevin Chen and Elisa DeStanchina (The Antitumor Assessment Core Facility). We thank the Molecular Cytology Core Facility and the Antitumor Assessment Core Facility in Memorial Sloan Kettering Cancer Center for help with experiments. Supplementary Fig. 2I, 5J and 5L is created with BioRender.com (https://biorender.com/). R.K. is supported by JSPS Overseas Research Fellowships.

## Author contributions

Conceptualization: SC, PR, RK, AS

Methodology: SC, PR, RK, AS, QL, HS, AF, EDS, JSRF, BW, SG

Visualization: RK, AS, HS, AF, EDS

Formal analysis: FA, AS, EDS, BW, SG

Investigation: RK, AS, QL, HS, EDS

Resources: HN, QJK

Supervision: SC, PR

Writing – original draft: SC, RK, AS, HS, EDS, AF

Writing – review & editing: SC, PR, AS, RK, QL, WM, AK, JSRF, BW, SG

Funding acquisition: SC, PR

## Data and materials availability

All data are available from the corresponding authors upon reasonable request.

