## Supplementary Figures for "Long-term breast cancer response to CDK4/6 inhibition defined by TP53-mediated geroconversion"

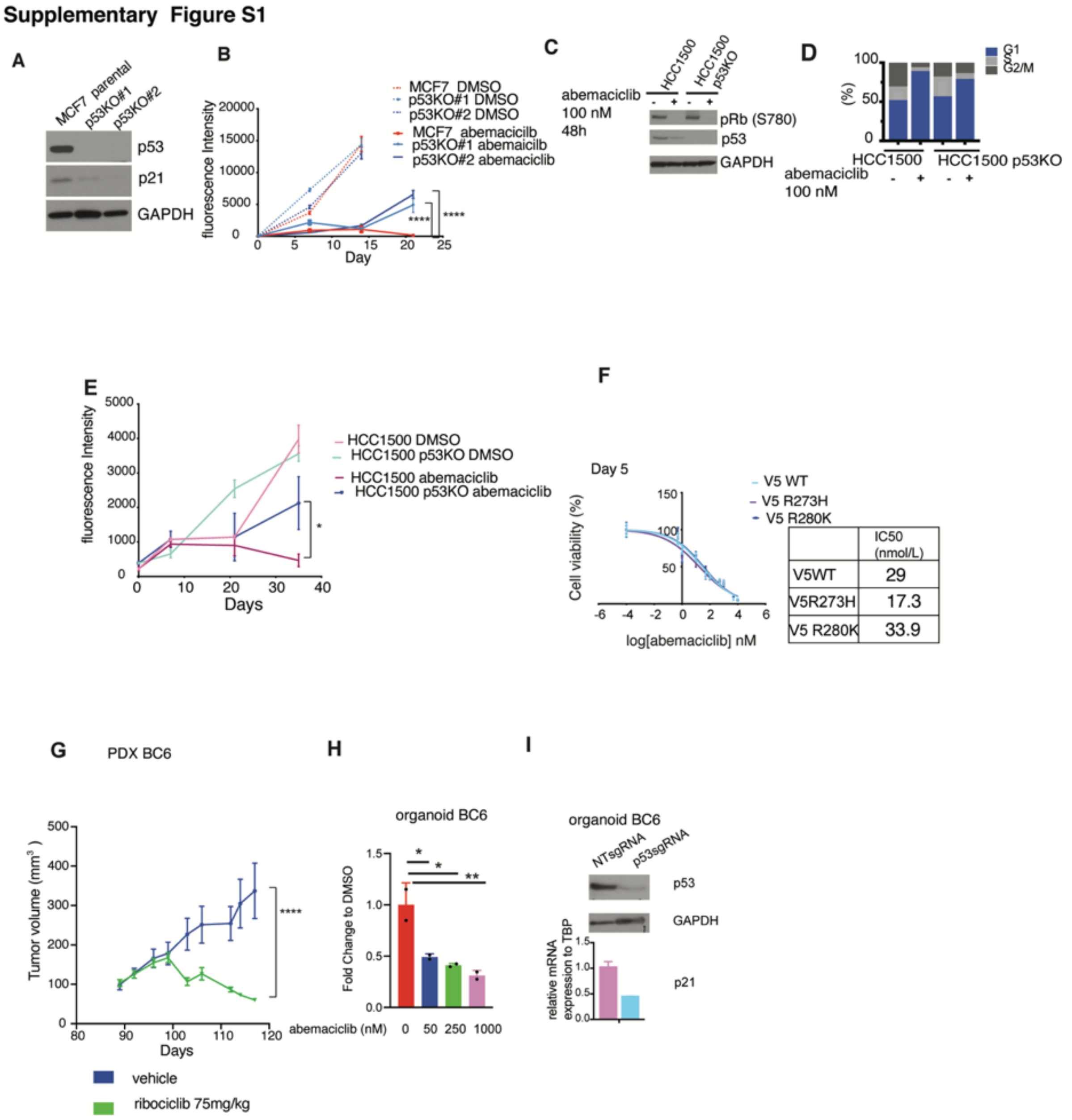


**Supplementary Figure S1. Loss of p53 led to regrowth after long-term culture.**

**A,** Immunoblotting of p53 and p21 in parental MCF7 and p53 knockout cell lines by two different sgRNA (p53KO#1 and p53KO#2). **B,** Cell viability of parental MCF7, p53KO clone 1 from p53KO#2 and p53KO clone 11 from p53KO#1 exposed to 50 nM abemaciclib or DMSO. Data are means of 4 biological replicates ± SEM. Two-way ANOVA, Tukey's multiple comparisons test. C**,** Immunoblotting of p53, p21 and phosphorylated Rb in parental HCC1500 cells and p53 knockout cells (HCC1500 p53KO) with 100 nM abemacilib for 48 h. **D**, Cell cycle distribution was measured after 48h 100 nM abemaciclib treatment in HCC1500 parental and p53KO cells. **E**, 5000 cells /well were seeded. Proliferation exposed to 500 nM abemaciclib or DMSO was analyzed. Data are means of 4 biological replicates ± SEM. Two-way ANOVA, Tukey's multiple comparisons test. **F,** Overexpression of V5-tagged wild type p53 (V5 WT), R273H (V5 R273H) and R280K mutants (V5 R280K) in p53 knockout cells. IC50s of abemaciclib were recorded at day4. **G,** Tumor volume change of PDX BC6 treated with vehicle or 75mg/kg ribociclib. Data are means of 3 biological replicates ± SEM. Two-way ANOVA Sidak's multiple comparison test. **H,** organoids BC6 were treated with indicated concertation of abemaciclib for 31 days. Bars are showing fold changes to DMSO. N=2 one way Anova, Dunnett's multiple comparisons test. **I,** Immunoblotting of p53 in organoid BC6 with NTsgRNA and p53sgRNA (top). Relative mRNA expression of p21 normalized to TBP. The ratio was represented as means ± s.d. n = 3 (bottom).

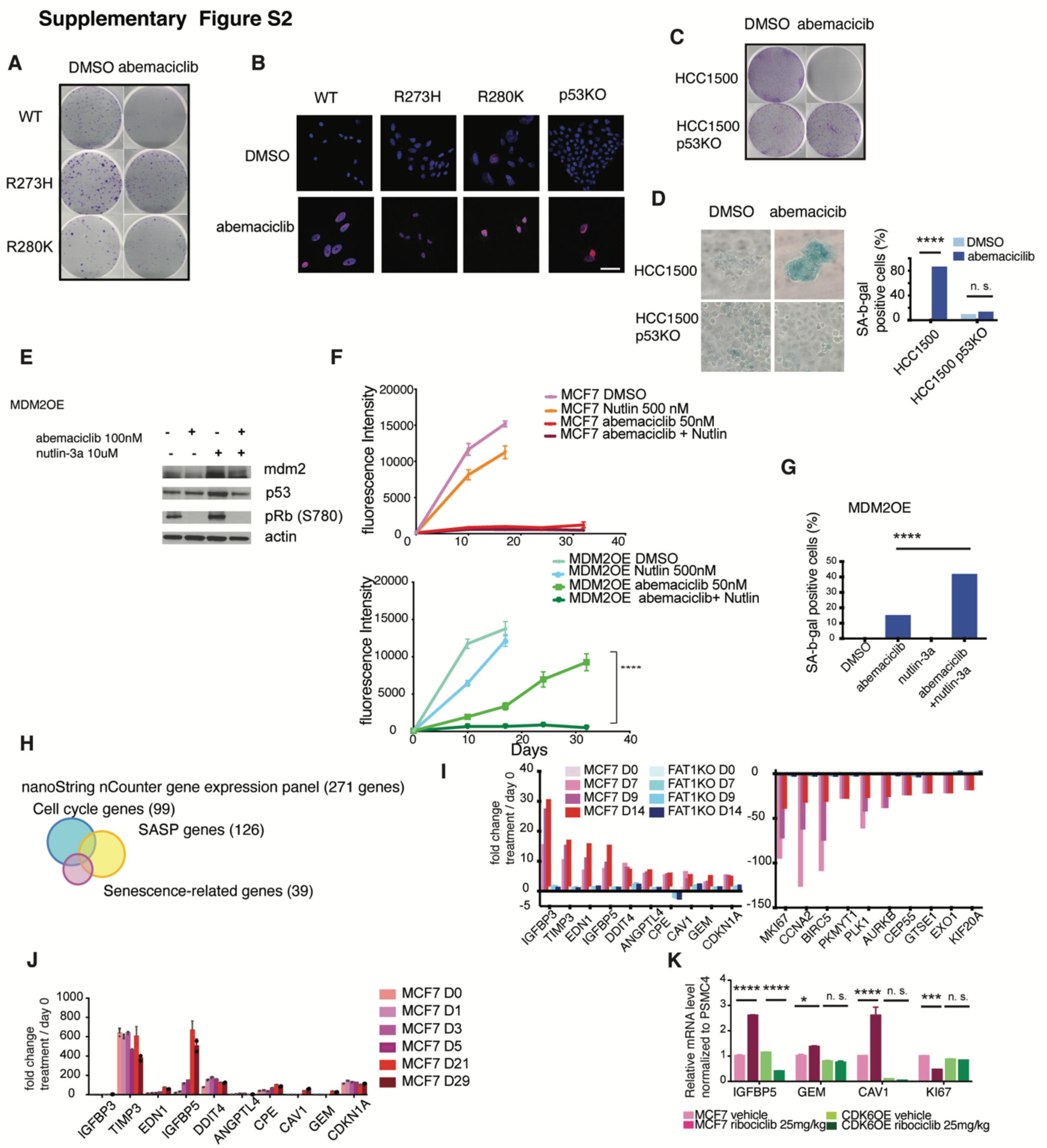


**Supplementary Figure S2. Lack of p53 prevents CDK4/6 inhibitor-induced senescence in HR$^{+}$ / wild type *TP53* breast cancer cell line**

A, Cells were treated with DMSO or 50 nM abemaciclib for 10 days, and 500 cells were replated after drug withdrawal for colony formation assay. **B,** V5-WT, V5-R273H, V5-R280K and p53KO cells were treated with 50 nM abemaciclib for 8 days. HP1γ foci and DAPI was determined by immunofluorescence. Representative images are shown. Scale bar, 50 μM. **C**, HCC1500 and HCC1500 p53KO cells were treated with DMSO or 100 nM abemaciclib for 10 days, and were reseeded after drug withdrawal for colony formation assay. **D**, Representative image of SA-β-gal activities in parental HCC1500 and HCC1500 p53KO with DMSO or 50 nM abemaciclib at day 7 (left). Bars are showing the percentage of SA-β-Gal positive cells (right). **E,** MDM2 overexpressing (MDM2 OE) cells were treated with DMSO, 100 nM abemaciclib or 1 μM Nutlin-3a for 48h. Immunoblotting shows indicated proteins. **F,** Cell viability of parental MCF7 and MDM2OE in response to DMSO, 50 nM abemaciclib, 250 nM Nutlin-3a and combination. Data are means of 4 biological replicates ± SEM. Two-way ANOVA, Tukey's multiple comparisons test. **G,** MDM2 OE cells were treated with DMSO, 100 nM abemaciclib, 500 nM Nutlin-3a or combination for 8 days. Bars are showing the percentage of SA-β-Gal positive cells..**H**, Custom nanoString nCounter gene expression panel contains 271 genes which are classified as cell cycle regulation genes, SASP genes and other senescence-related genes. **I,** The fold change of gene expression between day 0 and each indicated treatment days. MCF7 parental and FAT1KO cells were treated with 50 nM abemaciclib for indicated days. nCounter gene expression assays were performed for the Custom nanoString panel. The top 10 up-regulated or down-regulated genes in MCF7 parental cells on day 14 (red bar) compared to FAT1KO cells on day 14 (blue bar) were shown. The analysis was performed nSolver 4.0. Fold change is the same as indicated days/ day 0 when the value is greater than 1. When the value of indicated days/ day 0 is less than 1, then the fold change displays as the negative reciprocal, - indicated days/ day 0. **J**, The expression of senescence associated genes in RNA sequencing of MCF7 parental cells with 50 nM abemaciclib for indicated days. The fold change compared to the expression on day 0 was shown. Two-tailed unpaired Student's t tests (*p<0.05 in figure). **K,** qPCR of SASP expressions in MCF7 or CDK6OE xenograft tumors with 25 mg/kg ribociclib treatment. Relative mRNA expression of indicated genes normalized to PSMC4. The ratios are represented as means ± s.d. n = 3. Two-way ANOVA, Tukey's multiple comparisons test.

**Supplementary Figure S3**

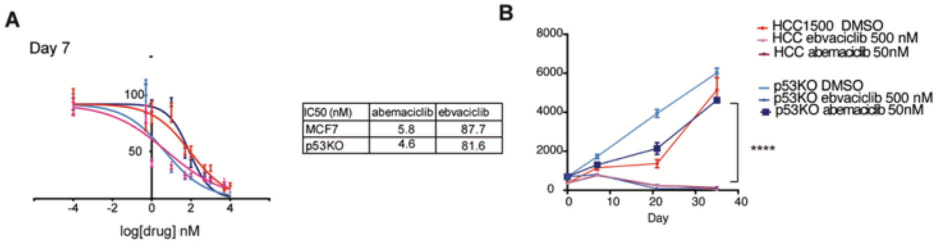


**Supplementary Figure S3. The combination of CDK2 and CDK4/6 inhibition in p53KO cells**

**A**, IC50s of abemaciclib or ebvaciclib in MCF7 parental and p53KO cells. **B,** Cell viability of HCC1500 parental and p53KO cells treated with DMSO, 500 nM ebvaciclib and 50 nM abemaciclib for indicated days. Data are means of 4 biological replicates ± SEM. One-way ANOVA followed by Tukey’s test.
